# Improved model quality assessment using sequence and structural information by enhanced deep neural networks

**DOI:** 10.1101/2022.08.12.503819

**Authors:** Jun Liu, Kailong Zhao, Guijun Zhang

## Abstract

Protein model quality assessment plays an important role in protein structure prediction, protein design, and drug discovery. In this work, DeepUMQA2, a substantially improved version of DeepUMQA for protein model quality assessment, is proposed. First, sequence features containing protein co-evolution information and structural features reflecting family information are extracted to complement model-dependent features. Second, a novel backbone network based on triangular multiplication update and axial attention mechanism is designed to enhance information exchange between inter-residue pairs. On CASP13 and CASP14 datasets, the performance of DeepUMQA2 increases by 20.5% and 20.4% compared with DeepUMQA, respectively (measured by top 1 loss). Moreover, on the three-month CAMEO dataset (March 11 to June 04, 2022), DeepUMQA2 outperforms DeepUMQA by 15.5% (measured by local AUC_0,0.2_) and ranks first among all competing server methods in CAMEO blind test. Experimental results show that DeepUMQA2 outperforms state-of-the-art model quality assessment methods, such as ProQ3D-LDDT, ModFOLD8, DeepAccNet, Atom_ProteinQA, and QMEAN3.

## 1 Introduction

Proteins are essential to life, and predicting their structures from sequences can facilitate the understanding of their functions. Various methods have been proposed to predict the three-dimensional structure of proteins [1-6]. The successful application of deep learning in recent years has significantly improved the accuracy of structure prediction [7-14], especially the end-to-end deep learning methods AlphaFold2 [15], whose prediction accuracy on most of the Critical Assessment of techniques for protein Structure Prediction (CASP14) targets is close to experimental results [16]. However, prediction accuracy for some proteins needs further improvement, and most prediction methods provide multiple prediction models. Therefore, it is important how to evaluate the accuracy of different prediction models without experimental structure. Estimating the accuracy of a protein model structure, or model quality assessment (MQA), is a crucial part of protein structure prediction and a gateway to the proper usage of models in biomedical applications [17]. Due to its importance, MQA has been a prediction category in CASP since 2006. In Continuous Automated Model EvaluatiOn (CAMEO), the performance of competing MQA servers is tested weekly by publishing prediction models of proteins whose experimental structures have not yet been disclosed.

MQA methods can be classified into single-model and multi-model methods according to the number of input models. Multi-model QA methods takes an ensemble of models as input and uses information from other models in the set to evaluate protein models, such as Pcons [18], APOLLO [19], ModFOLDclust [20], and QDistance [21]. Single-model QA methods evaluate a single model of inputs and does not depend on other models, such as the ProQ series [22-24], Ornate [25], GraphQA [26], DeepAccNet [27], and DeepUMQA [28]. In addition, there are a few quasi-single-model methods take a single model as an input and compare its similarity with a group of models that were built internally, such as MQAPsingle [29] and ModFOLD8 [30]. The performance of multi-model QA methods is limited by the number of input models and consensus of the models, whereas single-model QA methods are not subject to these limitations [31, 32]. Historically, multi-model methods are more accurate than single-model methods until CASP14. In CASP14, the best single-model QA method DeepAccNet [27] outperforms the best multi-model QA method, and the number of single-model methods accounts for more than 70% of all QA methods [17].

Current single-model quality assessment methods usually evaluate models through feature representation and neural network training, and the main differences between different methods are also in these two aspects. ProQ2 estimates model accuracy using a support vector machine (SVM) to predict the quality of a protein model by combing structural and sequence-based features calculated from the model [22]. ProQ3 uses Rosetta full-atom and coarse-grained centroid energy functions to descript the protein model, and trains the same SVM as ProQ2 to score the models [23]. ProQ3D show that a substantial improvement can be obtained using exactly the same inputs as in ProQ2 or ProQ3 but replacing the SVM by a deep neural network [24]. QAcon uses structural features, physicochemical properties and residue contact predictions as inputs to train a two-layer neural networks and evaluate models [31]. VoroMQA considers a protein structure as a set of balls corresponding to heavy atoms, and describes and integrates explicit interactions between atoms and the implicit interactions of protein atoms with solvents through contact areas [33]. Qdeep integrates distance information through stack depth ResNets to evaluate model quality [32]. QMEANDisCo employs the statistical potentials of mean force and consensus-based multi-template distance constraint score and trains a feed-forward neural network to adaptively trade off the contributions of both [34]. GraphQA [26] and ProteinGCN [35] represent the model as a graph and predict quality through graph convolutional networks. Ornate builds rotation- and translation-invariant 3D voxel atomic representations and predicts local and global model quality via deep 3D convolutional neural networks [25]. DeepAccNet uses 3D convolution and residual neural network to predict the inter-residue distance signed error and per-residue accuracy of the estimated model, which are further used as restraints to guide protein model refinement [27]. DeepUMQA introduces residue-level ultrafast shape recognition (USR) features to describe the topological relationship between residues and an overall structure, and combines with 1D, 2D and voxelization feature to assess the quality of the input model by a deep residual neural network [28].

In this work, DeepUMQA2, a significantly improved version of DeepUMQA, is proposed. On the basis of the features of the input model, sequence features from multiple sequence alignment and structural features from homologous templates are incorporated for the characterization of the potential properties of the model. Multiple sequence alignment (MSA) and homologous templates are first searched according to the sequence of the input model, and then sequence features and template structural features are extracted, and combined with input model-dependent features to form initial residue pair information. The pair information is updated iteratively through a new backbone network based on triangular multiplication update and axial attention mechanism. Then, two branch networks are used to predict the inter-residue distance deviation and the contact map with a threshold of 15Å, respectively, which are further used to calculate the accuracy of each residue of the model. The experimental results show that the performance of DeepUMQA2 has been greatly improved compared to DeepUMQA, and outperforms state-of-the-art methods. DeepUMQA2 ranks first in CAMEO’s blind test for three consecutive months (March 11 to June 04, 2022).

## 2 Methods

The pipeline of DeepUMQA2 is shown in Figure 1A. For the structure model to be evaluated, MSA and homologous templates are generated according to the sequence of the model by HHblits and HHsearch [36], respectively. Then, sequence features from MSA, structural features from the templates, and model-dependent features from the input model are extracted and concatenated into the initial pair features. The pair features are iteratively updated through a backbone network based on triangular multiplication update and axial attention mechanism and then through two independent residual networks for the prediction of inter-residue distance deviations and the contact map with a threshold of 15Å, respectively, which are further used in calculating the accuracy of each residue of the model.

**Figure 1.**
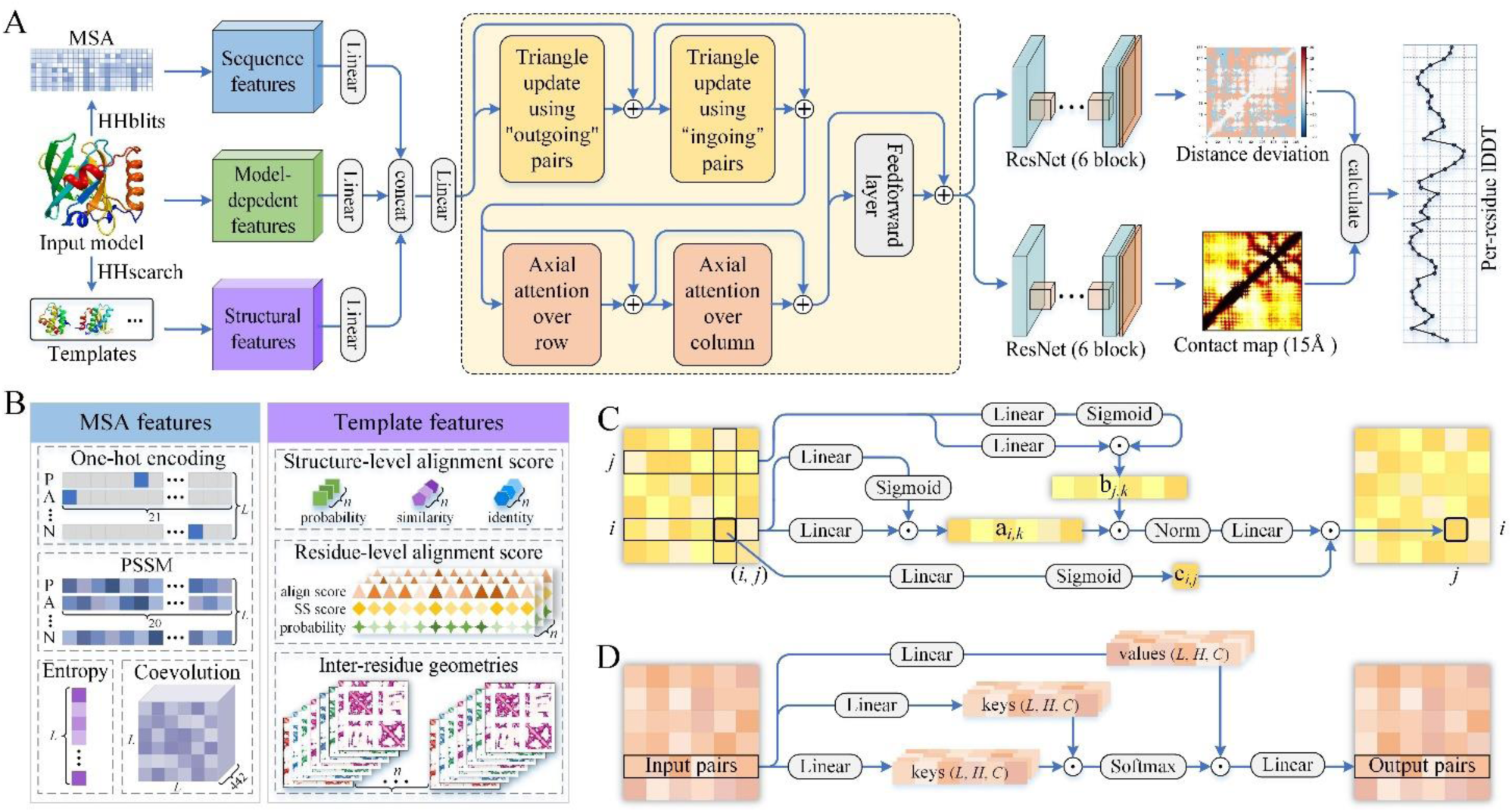
Overview of DeepUMQA2. (A) The pipeline of DeepUMQA2. (B) Sequence features extracted from MSA and structural features homologous templates. (C) Triangle update using “outgoing” pairs. (D) Axial attention over row.

### 2.1 Datasets

DeepUMQA2 is trained on the same dataset as DeepUMQA [28]. Briefly, a dataset was constructed from the PISCES server (deposited in May 2018) [37] and contains 7615 proteins with a maximum sequence redundancy of 40%. Each protein contained about 150 decoys generated by stochastic modeling, comparative modeling, deep learning modeling, and native structure perturbation. For the proteins in the training set, the sequence database used in searching the MSA and structural database for the search template were dated May 2018. In order to verify the performance of DeepUMQA2, we tested DeepUMQA2 on the model quality assessment datasets of CASP13, CASP14, and CAMEO, and compared with the top-performing server methods. The details of the CASP13, CASP14, and CAMEO datasets can be found in Supplementary Tables S1, S2, and S3.

The CASP13 dataset contained 76 protein targets, and each target had about 141 structural models submitted by server groups. Thus, 10739 models were obtained. The CASP14 dataset contained 70 protein targets, and each target had about 148 structural models. Thus, 10380 models were obtained. For the targets of CASP13 and CASP14, MSA was generated by searching the query sequence against the sequence database Uniclust30 (Version 2018_08), and homologous templates were generated by searching the structure database PDB100 (before May 01, 2018 for CASP13; before May 18, 2020 for CASP14). CAMEO dataset (March 11 to June 04, 2022) contained 192 protein targets, and each targets contained about 10 structural models submitted by the protein structure prediction servers, a total of 1882 proteins were obtained. For the target of CAMEO, MSA was generated by searching the query sequence against the sequence database UniRef30 (version 2022_02) [38] and BFD [39], and homologous templates were generated by searching the structure database PDB100 (before May 03, 2021).

### 2.2 Feature design

As shown in Figure 1A, the input features of DeepUMQA2 consisted of model-dependent features extracted from the input structure model, sequence features extracted from the MSA, and structural features from the homologous templates. The design of the model-dependent features was the same as DeeUMQA [28], including residue-level USR that characterize the relationship between residues and the topology structure, voxelization features that describe local information about each residue and its neighborhood, amino acid properties, secondary structure, Rosetta energy terms and inter-residue distance and orientations. Figure 1B clearly shows sequence and structural features.

#### 2.2.1 Sequence features from multiple sequence alignments

Sequences in MSA reflect the evolutionary information of a target protein and are thus useful in evaluating the quality of predicted models. For an evaluated model, the target sequence is first extracted, and then the MSA is generated by iteratively search against UniRef30 [38] and BFD [39] sequence databases using HHblits [36]. The E-value cutoff for sequence search is gradually relaxed (1e^−30^, 1e^−10^, 1e^−6^, and 1e^−3^) until the MSA has at least 2000 homologous sequences with 75% coverage or 5000 homologous sequences with 50% coverage. Sequence features extracted from MSA include 1D features: one-hot encoding of target sequence (*L* × 20, records the residue information of the target sequence, where *L* is the length of the model to be evaluated), residue position entropy (*L* × 1, records the positional information of residues), position-specific frequency matrix (*L* × 21, records the frequency of occurrence of 20 amino acids and one gap at each residue position in MSA), and 2D features: inverse covariance matrix and average product correction (*L* × *L* × 442, both used to describe coupling information between residues). Then, 1D features are extended to 2D features by horizontal and vertical striping and concatenated with 2D features.

#### 2.2.2 Structural features from homologous templates

Homologous templates contain direct structural constraint information, which can provide key information for model quality assessment. The homologous templates of the target sequence are detected by search against the PDB100 database with HHsearch [36]. A template is considered appropriate if its probability (the probability of template to be a true positive) is greater than 60 or its e-value is less than 0.001. The top 20 appropriate templates are selected for feature extraction. If the number of appropriate templates is less than 20, all good templates are used. If no appropriate template is available, template information is not used in model evaluation. As shown in Figure 1B, structural features extracted from templates has three levels. The first is residue pair level (2D features), including pairwise distances and orientations from template structures for aligned positions (*n* × *L* × *L* × 7, distance between C_β_ (C_α_ for GLY) and cosine and sine of orientations, where *n* is the number of templates). The second is residue level (1D features), including alignment probability, alignment confidence score, and secondary structure match score (*n* × *L* × 7). The third is the overall alignment level (scalar features), including homologous probability, sequence similarity, and sequence identity between the query and the template (*n* × 3). The 1D and scalar features are concatenated with 2D features by tiling them along both axes of 2D features.

### 2.3 Network architecture

As shown in Figure 1A, the network consists of three parts: an initial feature fusion to combine model-dependent features, sequence features and structural features; a backbone network based on triangular multiplication and axial attention mechanism to update pairs; a branch network based on residual blocks to predict the inter-residue distance deviation and the contact map. Features extracted from input model, MSA, and homologous templates are concatenated together to form an initial pair after passing through a linear layer. Two triangular multiplication update layers, two axial attention layers, and a feedforward layer constitute the pair update module, which are iterated three times. Triangular multiplication updates the residue pair representation by combining pair information within the triangle formed by current pair residues and other residues. Each residue pair (*i, j*) receives an update from the other pairs (*i, k*) and (*j, k*) of the triangles formed by residue *i, j* and the other residue *k*. Two symmetric versions: “outgoing” (using pair representation of *x*_*i,k*_ and *x*_*j,k*_ to update *x*_*i,j*_) and “ingoing” (using *x*_*k,i*_ and *x*_*k,j*_ to update *x*_*i,j*_) were obtained. Figure 1C shows the process of triangular multiplication update. The representation *x*_*i,j*_ of residue pair (*i, j*) is updated to *x′*_*i,j*_ by “outgoing” pairs according to the following formula:

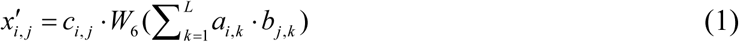

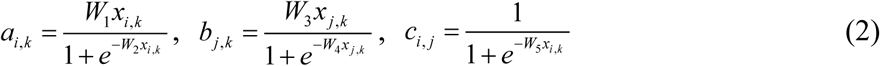

where *W*_1_ to *W*_6_ represent linear transformations. The triangular multiplication update is followed by two axial attention layers. Axial attention updates the residue pair representation along the rows and columns. Axial attention updates the residue pair (*i, j*) by combining information from all residue pairs that share residue *j* or *j*. Figure 1D shows the process of axial attention update. The representation *x*_*i,j*_ of residue pair (*i, j*) is updated to *x′*_*i,j*_ according to the following formula:

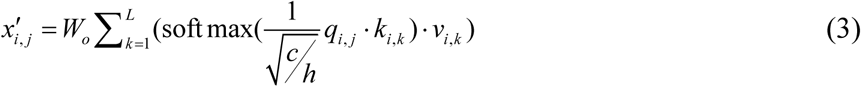

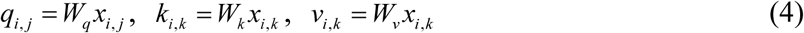

where *W*_*o*_, *W*_*q*_, *W*_*k*_, *W*_*v*_ represent linear transformations, *c* is the number of channels, and *h* is the number of heads. The feedforward layer consists of two linear layers and a GELU activation layer. The backbone network is followed by two branch networks to predict the inter-residue deviation and the contact map with a threshold of 15Å, respectively. Each branch network contains six residual blocks, and each residual block consists of two 2D convolutional layers (with dilation size 1, 2, 4, 8, 1 in turn), two instance normalization layers, two activation layers and a dropout layer.

### 2.4 Network model training

In this work, we represent the quality of a structure model by using the predicted per-residue lDDT, a superposition-free score for evaluating the local distance differences of atoms in a model [40]. To accurately predict the per-residue lDDT, we first predicted the inter-residue distance deviation and the contact map with a threshold of 15Å by using the neural network described above, and then we used them to calculate the lDDT of each residue. The inter-residue distance deviation was defined as the difference in the inter-residue C_β_ (C_α_ for GLY) distances between the input structure model and experimental structure and was divided into 15 bins with boundaries of ±0.5, ±1, ±2, ±4, ±10, ±15, and ±20Å. The predicted per-residue lDDT and global lDDT were calculated according to the following formulas:

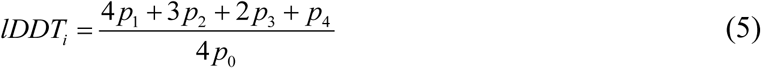

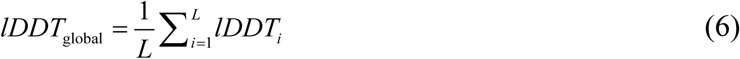

where *p*_0_ is the probability that the distance between the residue *i* and other residues is within 15 Å, and *p*_1_ is the probability that the absolute value of C_β_ distance deviation of all the residue pairs whose distance from the residue *i* within 15Å is less than 0.5 Å. In the same manner, *p*_2_, *p*_3_, and *p*_4_ are the probability that the absolute value of C_β_ distance deviation in all the residue pairs, whose distance from the residue *i* is within 15Å is 0.5Å–1.0Å, 1.0Å–2.0Å, and 2.0Å–4.0Å, respectively.

The loss of inter-residue distance deviation is evaluated by the multivariate cross-entropy loss function, the loss of contact is evaluated by the binary cross-entropy loss function, and the loss of per-residue lDDT is evaluated by the mean square loss function. The loss of the network consists of inter-residue distance deviation loss, contact loss and per-residue lDDT loss with weights of 1, 1 and 10. At each step of training, a single model from model sets of a randomly selected training protein. An epoch randomly traverses all training proteins, and a total of 100 epochs are trained. An AdamW optimizer with a “triangular” cycle learning rate (base_lr = 1e^−4^; max_lr = 5e^−4^; step_size_up = 10; step_size_down = 10) is used. The training and evaluation of the networks were performed using NVIDIA TITAN RTX GPUs.

## 3 Results

The performance of the model quality assessment is evaluated at global and local levels. Global evaluation metrics (Global QA) include the Pearson correlation coefficient, “top 1 loss”, area under the curve (AUC) of the receiver operating characteristic (ROC), the partial AUC of the ROC “trimmed” at a False Positive Rate (FPR) threshold of 0.2 (AUC_0,0.2_). The Pearson correlation coefficient shows the correlation between the predicted and real global scores of all protein models, and performance increases with correlation coefficient. “top 1 loss” is an important global evaluation metric in CASP [17], which is measured by using the absolute difference in lDDT between the best model selected by the predicted global score and the actual best model determined by the knowledge of the experimental structure. The lower the “top 1 loss”, the better the performance to find the best model. The AUC and AUC_0,0.2_ show the ability to distinguish good and poor models (or residues), which are calculated using lDDT score threshold of 0.6. Local evaluation metrics (Local QA) including Pearson correlation coefficient, Average residue-wise S-score (ASE), AUC, and AUC_0,0.2_. The roles of Person correlation coefficient, AUCs and AUC_0,0.2_ are the same as those of global evaluation, and the difference is that they are calculated on the local score. Noteably, CAMEO ranks MQA methods according to local AUC_0,0.2_. ASE measures the average residue-wise S-score error, and performance increases with ASE [17].

### 3.1 Performance on CASP13 and CASP14 datasets

The performance of DeepUMQA2 is tested on CASP13 and CASP14 datasets and compared with state-of-the-art methods, namely, ProQ2 [22], ProQ3 [23], ProQ3D [24], ProQ3D-lDDT [41], ProQ4 [42], VoroMQA [33], 3DCNN [43], ModFOLD7 [44], ModFOLD8 [45], GraphQA [26], QDeep [32], Ornate [25], QMEANDisCo [34], DeepAccNet [27] and DeepAccNet-MSA (A version of DeepAccNet incorporating distance predicted by trRosetta). The data of all the compared methods are obtained from the data archive on the official CASP website (https://predictioncenter.org/download_area/). The performance of DeepUMQA2 and comparison methods on CASP13 and CASP14 datasets are shown in Tables 1 and 2, respectively.

**Table 1.** Performance of DeepUMQA2 and comparison methods on CASP13 dataset. #Models represents the number of evaluated models.

| Method | Server ID | #Models | Global QA |  |  |  | Local QA |  |  |  |
| --- | --- | --- | --- | --- | --- | --- | --- | --- | --- | --- |
|  |  |  | Pearson | Top 1 Loss | AUC | AUC<sub>0,0.2</sub> | Pearson | ASE | AUC | AUC<sub>0,0.2</sub> |
| DeepUMQA2 | — | 10739 | 0.919 | 0.049 | 0.982 | 0.897 | 0.868 | 0.915 | 0.949 | 0.802 |
| ModFOLD7 | QA275_2 | 10739 | 0.826 | 0.076 | 0.897 | 0.706 | 0.736 | 0.827 | 0.883 | 0.650 |
| 3DCNN | QA359_2 | 7621 | 0.624 | 0.076 | 0.899 | 0.669 | 0.375 | 0.540 | 0.749 | 0.315 |
| ProQ2 | QA044_2 | 10739 | 0.715 | 0.070 | 0.887 | 0.649 | 0.632 | 0.752 | 0.851 | 0.530 |
| ProQ3 | QA187_2 | 10739 | 0.738 | 0.062 | 0.895 | 0.684 | 0.678 | 0.783 | 0.880 | 0.584 |
| ProQ3D | QA139_2 | 10739 | 0.801 | 0.071 | 0.908 | 0.713 | 0.700 | 0.807 | 0.869 | 0.566 |
| ProQ3D-IDDT | QA360_2 | 10739 | 0.850 | 0.065 | 0.933 | 0.770 | 0.782 | 0.896 | 0.902 | 0.657 |
| ProQ4 | QA440_2 | 10739 | 0.777 | 0.065 | 0.897 | 0.616 | 0.672 | 0.702 | 0.850 | 0.531 |
| VoroMQA-A | QA339_2 | 10739 | 0.672 | 0.078 | 0.857 | 0.596 | 0.599 | 0.737 | 0.819 | 0.427 |
| VoroMQA-B | QA030_2 | 10739 | 0.662 | 0.057 | 0.847 | 0.563 | 0.597 | 0.738 | 0.818 | 0.428 |

**Table 2.** Performance of DeepUMQA2 and comparison methods on CASP14 dataset. #Models represents the number of evaluated models.

| Method | Server ID | #Models | Global QA |  |  |  | Local QA |  |  |  |
| --- | --- | --- | --- | --- | --- | --- | --- | --- | --- | --- |
|  |  |  | Pearson | Top 1 Loss | AUC | AUC<sub>0,0.2</sub> | Pearson | ASE | AUC | AUC<sub>0,0.2</sub> |
| DeepUMQA2 | — | 10380 | 0.899 | 0.035 | 0.961 | 0.808 | 0.822 | 0.910 | 0.907 | 0.682 |
| DeepAccNet | QA403_2 | 10380 | 0.829 | 0.049 | 0.935 | 0.736 | 0.736 | 0.884 | 0.872 | 0.596 |
| DeepAccNet-MSA | QA209_2 | 10380 | 0.857 | 0.039 | 0.943 | 0.782 | 0.765 | 0.892 | 0.879 | 0.617 |
| ModFOLD8 | QA167_2 | 10380 | 0.629 | 0.080 | 0.835 | 0.454 | 0.572 | 0.804 | 0.786 | 0.435 |
| ProQ2 | QA067_2 | 9959 | 0.531 | 0.073 | 0.794 | 0.491 | 0.484 | 0.744 | 0.744 | 0.361 |
| ProQ3D | QA339_2 | 10380 | 0.651 | 0.068 | 0.846 | 0.512 | 0.575 | 0.788 | 0.786 | 0.420 |
| ProQ4 | QA309_2 | 9491 | 0.547 | 0.076 | 0.784 | 0.557 | 0.517 | 0.831 | 0.750 | 0.393 |
| QDeep | QA429_2 | 10380 | 0.656 | 0.087 | 0.858 | 0.544 | 0.499 | 0.767 | 0.745 | 0.345 |
| Ornate | QA346_2 | 10380 | 0.675 | 0.059 | 0.864 | 0.562 | 0.489 | 0.787 | 0.748 | 0.325 |
| GraphQA | QA210_2 | 9481 | 0.700 | 0.061 | 0.864 | 0.585 | 0.659 | 0.869 | 0.826 | 0.489 |
| QMEANDisCo | QA280_2 | 10380 | 0.767 | 0.083 | 0.874 | 0.629 | 0.671 | 0.871 | 0.829 | 0.513 |
| VoroMQA-dark | QA002_2 | 10380 | 0.565 | 0.062 | 0.787 | 0.449 | 0.547 | 0.746 | 0.783 | 0.389 |
| VoroMQA-light | QA102_2 | 10380 | 0.224 | 0.075 | 0.655 | 0.336 | 0.434 | 0.730 | 0.709 | 0.287 |

At the global evaluation level, DeeUMQA2 outperforms all the comparison methods on all performance metrics, whether on CASP13 or CASP14 datasets. The global Pearson correlation coefficients of DeepUMQA2 on the CASP13 and CASP14 datasets are 0.919 and 0.899, respectively, which are 8.1% and 4.9% higher than the second-best methods ProQ3D-lDDT and DeepAccNet-MSA, respectively. This result shows that DeepUMQA2 can capture the global accuracy of a model accurately. The “top 1 loss” of DeepUMQA2 on CASP13 and CASP14 datasets are 0.049 and 0.035, respectively, which are lower than all the comparison methods. The “top 1 loss” of DeepUMQA2 is 10.3% lower than that of DeepAccNet-MSA which achieves the lowest “top 1 loss” in CASP14. This result shows that our method is better in identifying the best model from a large number of predictive models. To analyze the ability of DeepUMQA2 and the comparison methods to distinguish between good and poor models, we performed ROC analysis based on the predicted global accuracy versus the actual global accuracy of all models. Figures 2(A) and (B) show the ROC curves of different methods on the CASP13 and CASP14 datasets, respectively. The AUCs of DeepUMQA2 on the CASP13 and CASP14 datasets are 0.982 and 0.961, respectively, and are significantly better than comparison methods. The AUC_0,0.2_ of DeepUMQA2 outperforms all comparison methods on the CASP13 and CASP14 datasets.

**Figure 2.**
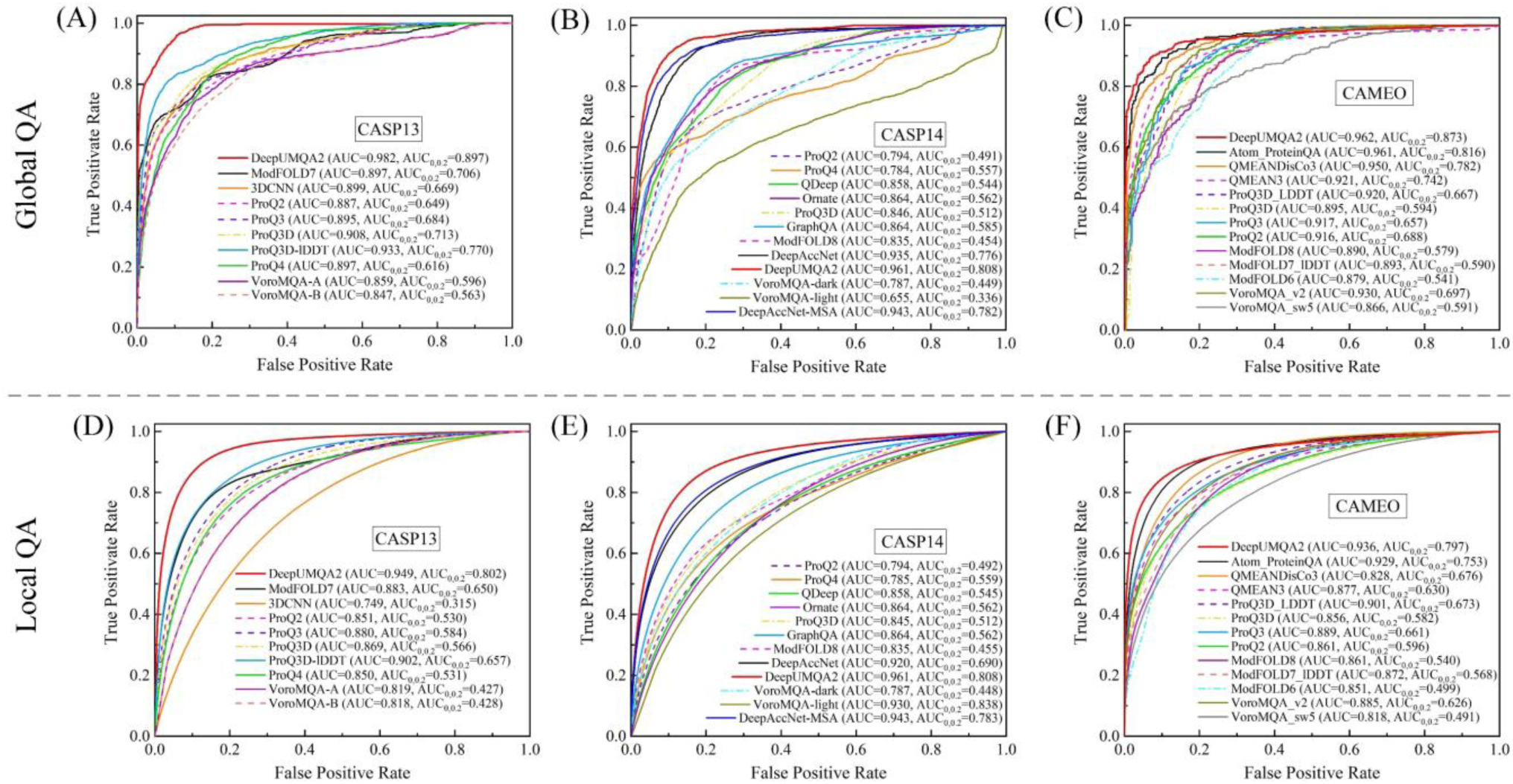
Receiver operating characteristic (ROC) curve of DeepUMQA2 and the comparison model quality assessment methods in global evaluation level (Global QA) on (A) CASP13, (B) CASP14, and (C) CAMEO datasets; in local evaluation level (Local QA) on (D) CASP13, (E) CASP14, and (F) CAMEO datasets. An lDDT cutoff value of 0.6 is used in distinguishing between good and poor models in Global QA (residues in Local QA).

At the local evaluation level, DeepUMQA2 still has the best performance among all methods, on the CASP13 and CASP14 datasets. The local Pearson correlation coefficients of DeepUMQA2 on the CASP13 and CASP14 datasets are 0.868 and 0.822, respectively, which are about 11.0% and 7.5% higher, respectively, than those of the second-best methods ProQ3D-lDDT and DeepAccNet-MSA. The ASE of DeepUMQA2 is 0.915 and 0.910 on the CASP13 and CASP14 datasets, respectively, significantly outperforming all comparison methods. This result shows that the per-residue accuracy predicted by our method is more precise overall. We also plot ROC curves based on predicted and actual accuracy for all the residues of models. As shown in Figures 2(D) and (E), DeepUMQA2 has the highest AUC and AUC_0,0.2_ among all methods on the CASP13 and CASP14 datasets. This result suggests that the local accuracy score predicted by DeepUMQA2 can distinguish between accurate and inaccurate residues in the structure model.

To further analyze the combined performance of DeepUMQA2 and comparison methods, we ranked these methods using the sum of the Z-scores of all global performance metrics and all local performance metrics, respectively. Figure 3 shows the comprehensive performance of all mthods in the global and local accuracy estimation. It can be found that the global and local comprehensive performance of DeepUMQA2 is the best among all methods on the CASP13 and CASP14 datasets.

**Figure 3.**
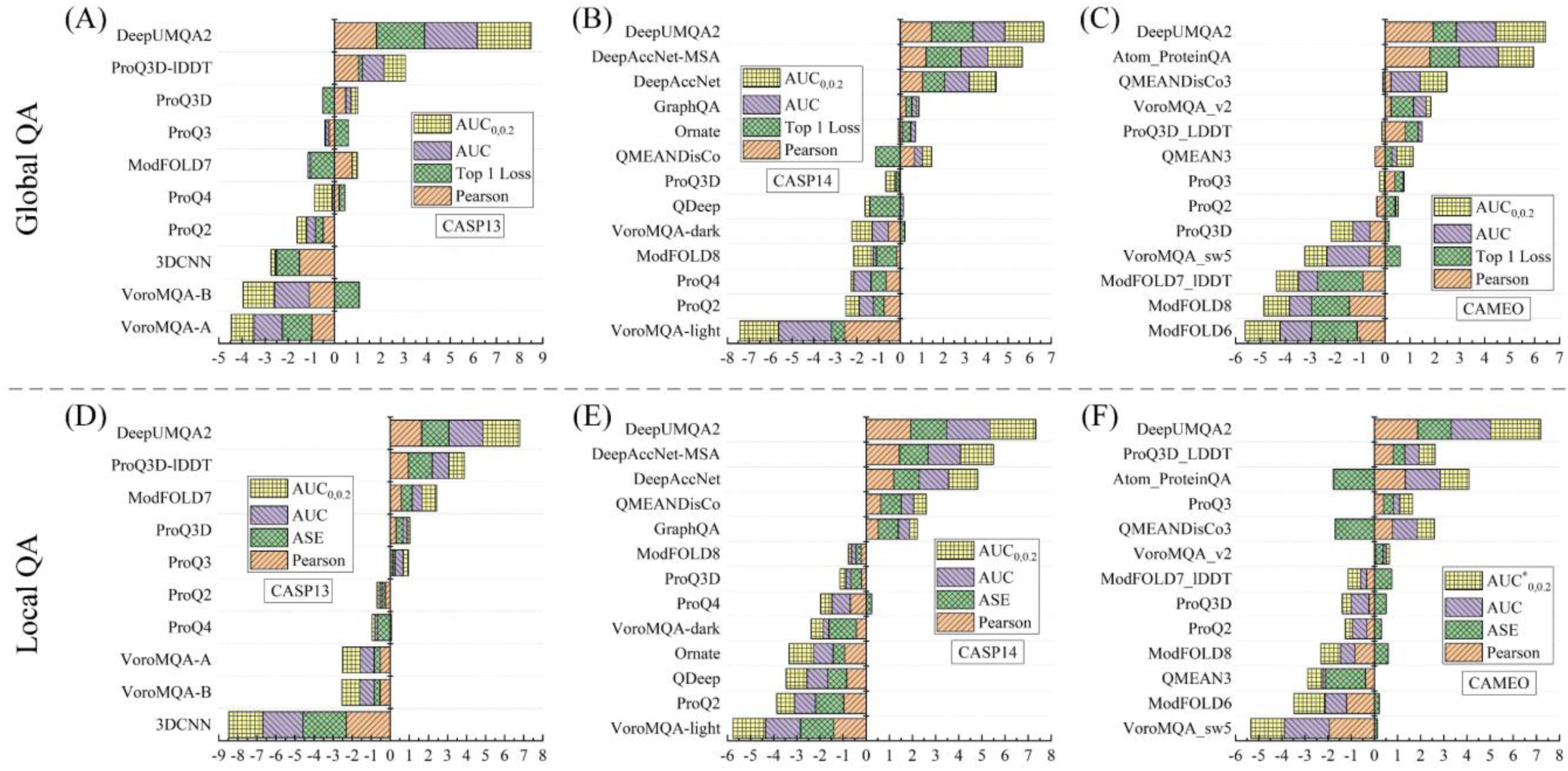
Ranking of the methods in the global and local accuracy evaluation (Global QA and Local QA). (A), (B), and (C) are ranked according to the sum of the Z-scores of all the Global QA metrics on the CASP13, CASP14, and CAMEO datasets, respectively. (D), (E), and (F) are ranked according to the sum of the Z-scores of all the Local QA metrics on the CASP13, CASP14, and CAMEO datasets, respectively.

Note that the 3DCNN in Table 1 only contains 7621 models, and all other methods have 10739 models. Some models are also missing from ProQ2, ProQ4 and GraphQA in Table 2, and the number of models evaluated by other methods is 10380. The comparison of DeepUMQA2 with these methods applied to the same number of models is shown in Supplementary Tables S4 and S5. The results show that DeepUMQA2 still outperforms these comparison methods.

### 3.2 Performance on CAMEO datasets

To further validate the performance of DeepUMQA2, we participated in the blind test of CAMEO-QE (https://www.cameo3d.org/quality-estimation/3-months/quality/all/) for three consecutive months (March 11 to June 04, 2022), and ranked first among all participating MQA methods (see Supplementary Figure S1). In this section, we analyzed the performance of DeepUMQA2 and other top-performing participating server methods in detail on three month CAMEO dataset. All model evaluation data were obtained from the CAMEO official website. Comparison methods include Atom_ProteinQA, QMEAN3 [46], QMEANDisCo3 [47], ProQ2 [22], ProQ3 [23], ProQ3D [24], ProQ3D_LDDT [41], ModFOLD6 [48], ModFOLD7_LDDT [44], ModFOLD8 [45], VoroMQA_v2, and VoroQA_sw5 [33].

The performance of all methods on global and local metrics is shown in Table 3. At the global evaluation level, the Pearson correlation coefficient of DeepUMQA2 is 0.899, which is superior to other methods. The top 1 loss of DeepUMQA2 is 0.017, second only to that of Atom_ProteinQA. The global AUC and AUC_0,0.2_ of DeepUMQA2 are the highest. The AUC_0,0.2_ of DeepUMQA2 is significantly higher than that of other methods, and is about 7.0% higher than that of the second-best method Atom_ProteinQA. At the local evaluation level, DeepUMQA2 outperforms all other methods on all performance metrics. The Pearson correlation coefficient and ASE of DeepUMQA2 are 0.870 and 0.910, respectively, which are about 4.9% and 6.1% higher, respectively, than those of the second-best Atom_ProteinQA (Perason: 0.829) and ModFOLD7_lDDT (ASE: 0.858). It is worth noting that the local AUC_0,0.2_ of DeepUMQA2 is 0.797, which is significantly higher than other methods and about 10.8% higher than the suboptimal method Atom_ProteinQA. Noted that CAMEO officially ranks MQA methods with a local AUC_0,0.2_. We also rank all methods by using the sum of the Z-score all global evaluation metrics (Figure 3C) and local evaluation metrics (Figure 3F), respectively, to analyze comprehensive performance. It can be found that DeepUMQA2 achieves the best comprehensive performance in both Global QA and Local QA.

**Table 3.** Performance of DeepUMQA2 and top-performance sever methods on the CAMEO dataset (March 11 to June 04, 2022). #Models represents the number of evaluated models.

| Server name | Server ID | #Models | Global QA |  |  |  | Local QA |  |  |  |
| --- | --- | --- | --- | --- | --- | --- | --- | --- | --- | --- |
|  |  |  | Pearson | Top 1 Loss | AUC | AUC<sub>0,0.2</sub> | Pearson | ASE | AUC | AUC<sub>0,0.2</sub> |
| DeepUMQA2 | server43 | 1860 | 0.899 | 0.017 | 0.962 | 0.873 | 0.870 | 0.910 | 0.936 | 0.797 |
| Atom_ProteinQA | server41 | 1313 | 0.891 | 0.008 | 0.961 | 0.816 | 0.829 | 0.675 | 0.929 | 0.719 |
| QMEANDisCo3 | server29 | 1881 | 0.800 | 0.057 | 0.950 | 0.782 | 0.786 | 0.681 | 0.915 | 0.676 |
| QMEAN3 | server20 | 1882 | 0.763 | 0.043 | 0.921 | 0.742 | 0.695 | 0.678 | 0.877 | 0.558 |
| ProQ3D_LDDT | server31 | 1302 | 0.834 | 0.034 | 0.920 | 0.667 | 0.788 | 0.841 | 0.901 | 0.673 |
| ProQ3D | server33 | 1296 | 0.750 | 0.047 | 0.895 | 0.594 | 0.708 | 0.841 | 0.856 | 0.582 |
| ProQ3 | server32 | 1301 | 0.809 | 0.042 | 0.917 | 0.657 | 0.756 | 0.835 | 0.889 | 0.661 |
| ProQ2 | server8 | 1873 | 0.767 | 0.038 | 0.916 | 0.688 | 0.700 | 0.826 | 0.861 | 0.586 |
| ModFOLD8 | server39 | 1680 | 0.702 | 0.114 | 0.890 | 0.579 | 0.661 | 0.847 | 0.861 | 0.540 |
| ModFOLD7_IDDT | server28 | 1458 | 0.734 | 0.125 | 0.893 | 0.590 | 0.700 | 0.858 | 0.872 | 0.568 |
| ModFOLD6 | server21 | 1723 | 0.721 | 0.126 | 0.879 | 0.541 | 0.633 | 0.819 | 0.851 | 0.499 |
| VoroMQA_v2 | server17 | 1882 | 0.800 | 0.018 | 0.930 | 0.697 | 0.730 | 0.827 | 0.885 | 0.626 |
| VoroMQA_sw5 | server15 | 1882 | 0.749 | 0.030 | 0.866 | 0.591 | 0.576 | 0.813 | 0.818 | 0.491 |

It should note that the number of models evaluated by different server methods may be inconsistent, as some server methods may miss some models when submitting results. The data in Table 3 is based on the actual prediction model of each server method (the same as CAMEO’s official practice). For in-deep analysis, we compared DeepUMQA2 with each method separately to ensure that the number of models for comparison is the same. The comparison results are shown in Supplementary Table S6. It can be found that the overall performance of DeepUMQA2 is still better than that of all other methods.

### 3.3 Ablation test

This section analyzes the impact of sequence features, structural features, and attention-based network architecture on DeepUMQA2 performance. We trained two comparison versions using the same data set and parameters: DeepUMQA2 (w/o sequence) without sequence features and DeeoUMQA2 (w/o sequence & structure) without sequence and structural features. DeepUMQA2 (w/o sequence & structure) uses only the structural information of the model for evaluation, similar to DeepUMQA, except that they use different network architectures. The backbone network of DeepUMQA uses stacked residual blocks, whereas DeepUMQA2 (w/o sequence & structure) uses triangular multiplication update and axial attention network. We compared the performance of DeeUMQA2, DeeUMQA2 (w/o structure), DeeUMQA2 (w/o sequence & structure) and DeepUMQA on CASP13, CASP14, and CAMEO datasets.

As shown in Table 4, DeepUMQA2 (w/o sequence & structure) outperforms DeepUMQA in all global performance metrics on the three test sets. Both are comparable in terms of local ASE and AUC, whereas DeepUMQA2 (w/o sequence & structure) outperforms DeepUMQA in terms of local Pearson correlation coefficient and AUC_0,0.2_. The sum of Z-scores of all the global or local evaluation metrics in Figure 4 shows that DeepUMQA2 (w/o sequence & structure) outperforms DeepUMQA in global and local comprehensive performance on the three test sets. The possible reason is that triangular multiplication update not only enhances the pair update but also alleviates information conflict among residual pairs to a certain extent. The multi-head axial attention mechanism may capture pairing information at a deeper level. On the CASP13 and CASP14 datasets, DeepUMQA2 (w/o structure) outperforms DeepUMQA2 (w/o sequence & structure) on all global and local performance metrics, and DeepUQA2 outperforms DeepUMQA2 (w/o structure) across the board. On the CAMEO dataset, DeepUMQA2 outperforms DeepUMQA2 (w/o structure) on all local performance metrics, and DeepUMQA2 (w/o structure) outperforms DeepUMQA2 (w/o sequence & structure). This trend also appears in global performance metrics except “top 1 loss”. In contrast to CASP13 and CASP14 datasets, the “top 1 loss” of DeepUMQA2 on CAMEO datasets increases gradually with the gradual addition of sequence and structural features. This increase may be due to the low number of models per target protein in CAMEO, averaging less than 10. Note that CAMEO officially ranks the MQA methods according to the local AUC_0,0.2_. The sum of Z-scores of all performance metrics in Figure 4 shows that the global comprehensive performance and local comprehensive performance of DeepUMQA2 gradually and significantly improves after the gradual addition of sequence features and structural features on the CASP13, CASP14, and CAMEO datasets.

**Table 4.** Performance of the different versions of DeepUMQA2 and DeepUMQA on CASP13, CASP14, and CAMEO datasets.

| Dataset | Method | Global QA |  |  |  | Local QA |  |  |  |
| --- | --- | --- | --- | --- | --- | --- | --- | --- | --- |
|  |  | Pearson | Top 1 Loss | AUC | AUC<sub>0,0.2</sub> | Pearson | ASE | AUC | AUC<sub>0,0.2</sub> |
| CASP13 | DeepUMQA2 | 0.919 | 0.049 | 0.982 | 0.897 | 0.868 | 0.915 | 0.949 | 0.802 |
|  | DeepUMQA2(w/o structure) | 0.890 | 0.050 | 0.947 | 0.803 | 0.817 | 0.903 | 0.914 | 0.711 |
|  | DeepUMQA2(w/o sequence & structure) | 0.838 | 0.052 | 0.944 | 0.778 | 0.762 | 0.891 | 0.900 | 0.651 |
|  | DeepUMQA | 0.837 | 0.059 | 0.939 | 0.766 | 0.745 | 0.892 | 0.900 | 0.644 |
| CASP14 | DeepUMQA2 | 0.899 | 0.035 | 0.961 | 0.808 | 0.822 | 0.910 | 0.907 | 0.682 |
|  | DeepUMQA2(w/o structure) | 0.836 | 0.040 | 0.932 | 0.770 | 0.728 | 0.895 | 0.858 | 0.596 |
|  | DeepUMQA2(w/o sequence & structure) | 0.807 | 0.041 | 0.924 | 0.701 | 0.713 | 0.890 | 0.855 | 0.559 |
|  | DeepUMQA | 0.799 | 0.044 | 0.920 | 0.690 | 0.710 | 0.890 | 0.856 | 0.553 |
| CAMEO | DeepUMQA2 | 0.899 | 0.017 | 0.962 | 0.873 | 0.870 | 0.910 | 0.936 | 0.797 |
|  | DeepUMQA2(w/o structure) | 0.882 | 0.012 | 0.958 | 0.838 | 0.859 | 0.904 | 0.933 | 0.775 |
|  | DeepUMQA2(w/o sequence & structure) | 0.877 | 0.010 | 0.955 | 0.834 | 0.845 | 0.898 | 0.924 | 0.746 |
|  | DeepUMQA | 0.872 | 0.011 | 0.951 | 0.833 | 0.842 | 0.898 | 0.925 | 0.690 |

**Figure 4.**
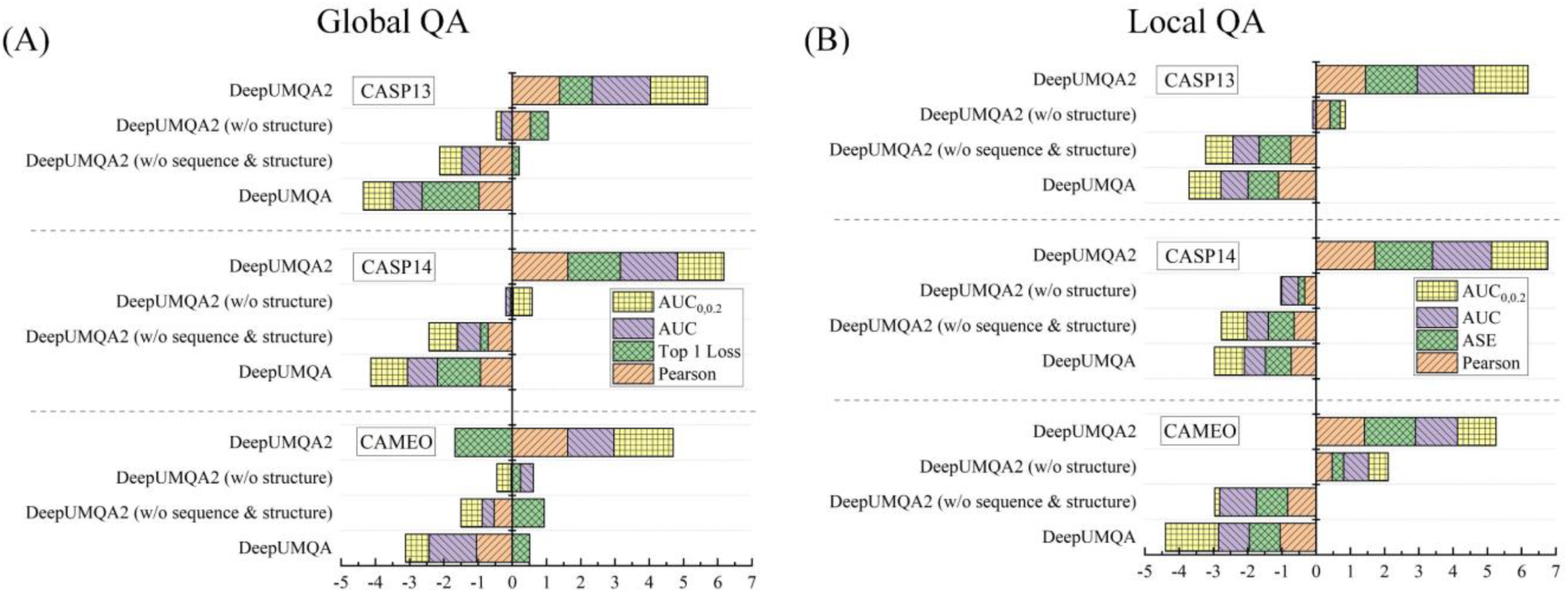
Ranking of different versions of DeepUMQA2 and DeepUMQA in the global and local accuracy evaluation (Global QA and Local QA). (A) Rankings based on the sum of the Z-scores of Global QA on the CASP13, CASP14, and CAMEO datasets. (B) Rankings based on the sum of the Z-scores of Local QA on the CASP13, CASP14, and CAMEO datasets.

### 3.4 Case study

This section quantitatively analyzes the quality assessment of prediction models for individual proteins. Figure 5 shows quantitative analysis on the CASP14 target T1034, which contains 145 prediction models and the real global lDDT is distributed in [0.582, 0.752]. Figure 5(B) shows the head-to-head comparison of the real global lDDT and DeepUMQA2 predicted global lDDT on these 145 models, which are strongly correlated. The Pearson correlation coefficient is 0.917, the R-square value is 0.841, and the best model selected according to the predicted global lDDT is the actual best model. Figure 5(C) plots the global and local ROC curves, the global AUC and AUC_0,0.2_ are both 0.993, the local AUC and AUC_0,0.2_ are 0.947 and 0.756, respectively, and the ASE is 0.945. Figures 5(D) to (F) present the structures of the three models with different precision, their corresponding predicted and actual global lDDTs, and per-residue predicted and real lDDT scores. It can be found that the per-residue lDDT predicted by DeepUMQA2 can capture the trend of residue accuracy variation, and it is easy to distinguish between high- and low-precision residue regions, which can provide favorable information for further model refinement.

**Figure 5.**
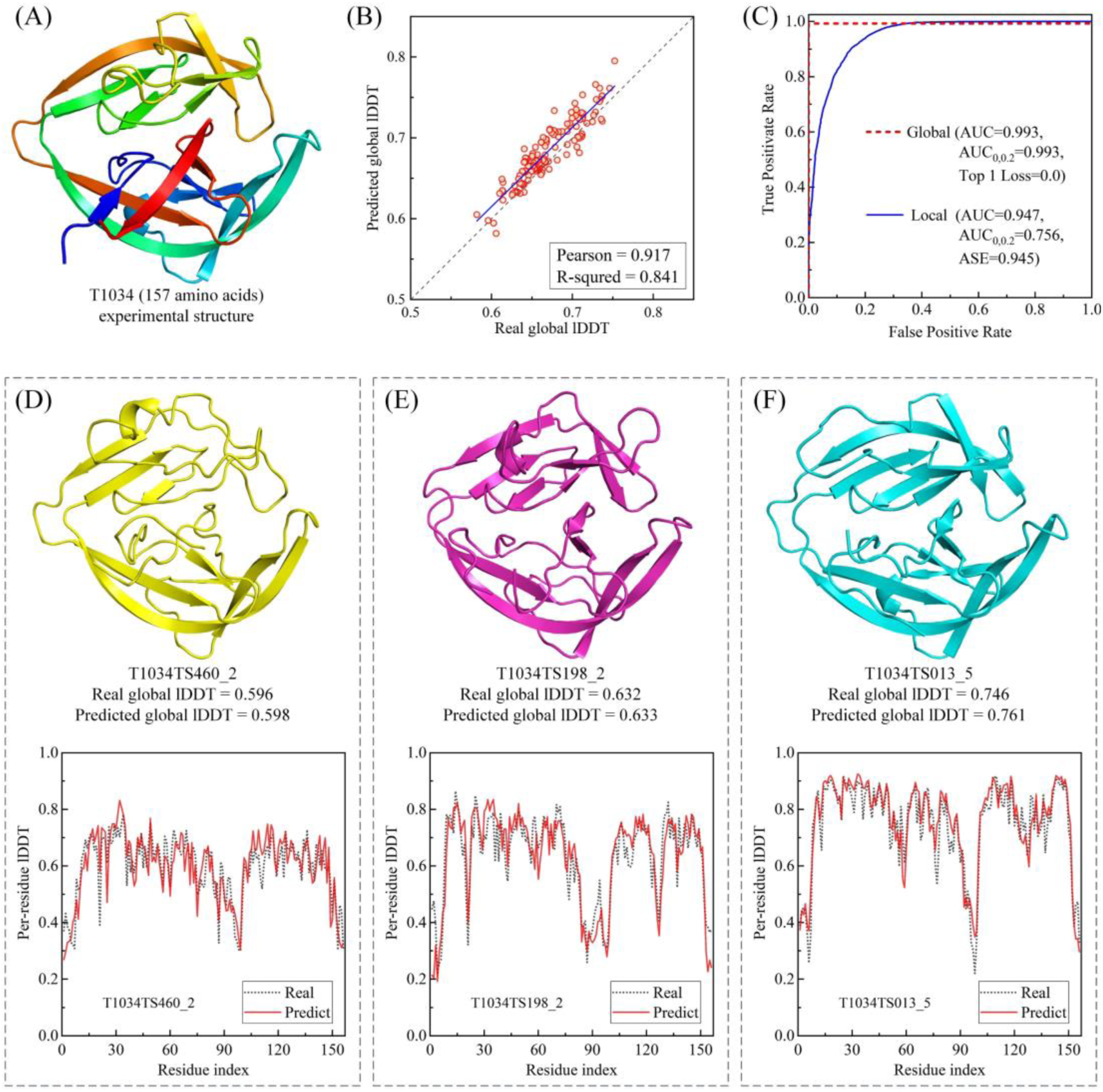
Qualitative analysis of the performance of DeepUMQA2 on target T1034 of CASP14. (A) The experimental structure of T1034. (B) Head-to-head comparison of the predicted lDDT and the real lDDT for all the structure models of T1034. (C) The global and local ROC curve of DeepUMQA2. (D) to (E) Results of the model assessment of three randomly selected models of target T1034 with different quality.

## 4 Conclusion

We propose DeepUMQA2, a significantly improved version of DeepUMQA. The major improvements come from two aspects. On the one hand, sequence and structural features are integrated on the basis of the model-dependent features to describe the co-evolution and family information of protein models comprehensively. On the other hand, the network backbone adopts a novel network based on triangular multiplication update and axial attention mechanism to replace the original residual network. Extensive experimental results show that the performance of DeepUMQA2 is significantly improved and outperforms state-of-the-art model quality assessment methods. DeepUMQA2 ranks first in CAMEO’s model quality evaluation in blind tests for three months (March 11 to June 04, 2022). Significant improvement in performance comes from two main sources. On the one hand, sequence features extracted from MSA and structural features extracted from templates contain protein co-evolution and family information, which may provide key protein-level information for assessing model quality and complement the inherent information of the model to be evaluated. On the other hand, network architecture based on triangular multiplication updates and axial attention may be able to better implement residual pair information exchange. Significant breakthroughs have been made in the prediction accuracy of single-chain protein structures, but there is still a lot of room for improvement in the prediction accuracy of complex structures. How to accurately assess the quality of the complex structures and improve the prediction accuracy of a complex structures is the direction that needs to be undertaken in the future.

## Data and code availability

All data needed to evaluate the conclusions are present in the paper and the Supplementary Materials. The web server of DeepUMQA2 is freely available at http://zhanglab-bioinf.com/DeepUMQA2.

